# The grapevine ABC transporter B family member 15 (VvABCB15) is trans-resveratrol transporter out of grapevine cells

**DOI:** 10.1101/2023.10.20.563313

**Authors:** A Martínez-Márquez, V Martins, S Sellés-Marchart, H Gerós, P Corchete, R Bru-Martínez

**Affiliations:** Plant Proteomics and Functional Genomics Group, Department of Agrochemistry and Biochemistry, Faculty of Science, University of Alicante, Alicante, Spain; Center for the Research and Technology of Agro-Environmental and Biological Sciences (CITAB), Quinta de Prados, Vila Real 5001-801, Portugal; Research Group in Applied Plant Biology and Innovation in Agrofood-Agrobioplant, Department of Biology, School of Sciences, University of Minho, Campus de Gualtar, Braga 4710-057, Portugal; Research Technical Facility, Proteomics and Genomics Division, University of Alicante, Alicante, Spain; Department of Plant Physiology, Campus Miguel de Unamuno, University of Salamanca, E-37007 Salamanca, Spain

**Keywords:** ATP binding cassette -ABC- transporter B family member 15, Label-free proteomics analysis, Grapevine cell culture, Resveratrol, Transformation, Transport, *Vitis vinifera*

## Abstract

Stilbenes, particularly *trans*-resveratrol, play a highly relevant defense role in grapevine as phytoalexin induced in response to stress. Metabolism and transport of stilbenes can be conveniently investigated in grapevine cell culture since large amounts of *trans*-resveratrol are accumulated in the extracellular medium upon treatment with the elicitor methylated cyclodextrin, either alone or combined with methyl jasmonate. Aiming at finding trans-resveratrol transporter candidates a proteomic approach on grapevine cells membrane fractions was performed. The candidate VvABCB15 was functionally characterized. Its stable expression in both yeast and *Silybum marianum* cells heterologous systems led to increased *trans*-resveratrol transport in these hosts. Transient expression in *Vitis* cells showed an enhanced absorbent- or elicitor-assisted accumulation of extracellular *trans*-resveratrol in both VvABCB15-expressing or VvGSTU10/VvABCB15-coexpressing cell suspension cultures. Experiments of transient expression in *Vitis* cell suspensions using light-switchable stilbene synthase (pHYH::VvSTS3) and VvABCB15 further confirmed the role of the candidate as *trans*-resveratrol transporter. VvABCB15-YFP fusion proteins in *Nicotiana* leaf showed localization in plasma membrane, being consistent with a functional role in *trans*-resveratrol transport. This is the first report providing evidence for the involvement of an ABC transporter B-type, VvABCB15 in *trans*-resveratrol transport to the extracellular medium of grapevine cells.

## INTRODUCTION

Plants are permanently challenged by both abiotic and biotic stresses and to cope with them, plants endow a complex defense and resistance network of constitutive and inducible mechanisms, including phytoalexins. Stilbenes are the phytoalexin group found in grapevine (*Vitis vinifera*) (Langcake and Pryce, 1977), in Vitaceae and few other plant families (Morales et al. 2000). The biosynthetic route leading to stilbenes is a small branch of the general phenylpropanoid pathway consisting of the condensation of the precursor p-coumaroyl-CoA with 3 units of malonyl-CoA catalyzed by stilbene synthase (STS) (Langcake and Pryce, 1977) to produce *trans*-resveratrol (3,4′,5-trihydroxystilbene, *t*-R), which is the most abundant stilbene both in grapevine vegetative tissues and in berries as well as in cultured cells in response to abiotic and biotic stress (Cantos et al., 2003; Wang et al., 2010). This primary stilbene has phytoalexin activity by itself but it can be further metabolized to be converted into more active stilbene phytoalexins such as the methylated derivative pterostilbene (Adrian et al. 1997) by grapevine resveratrol O-methyltransferase (Schmidlin et al. 2008). Furthermore, fungal laccases (Pezet, 1998; Schouten et al. 2002) can generate the dimer ε−Viniferin and higher oligomers (Langcake, 1981; Gabaston et al. 2017) having stronger antifungal activity than *t*-R, thus emphasizing the relevance of the extracellular *t*-R metabolism in its role as phytoalexin.

In addition to function as phytoalexins, a strong focus on stilbenes bioproduction research has been implemented for the last decade (Jeandet et al. 2021) due to a large and diverse number of biological activities attributed mainly to *t*-R, including inhibiting the progression of cardiovascular, carcinogenic and neurodegenerative diseases as well as the ageing process, as confirmed by several *in vitro* assays (Kukreja et al., 2014).

Treatment of grapevine cell culture with either abiotic or biotic elicitors promotes the expression of STS genes and proteins followed by accumulation of intra- and extracellular resveratrol (Tassoni et al. 2005; Lijavetzky et al. 2008; Martinez-Esteso et al. 2009; Almagro et al., 2014). The extracellular accumulation is particularly abundant when methylated cyclodextrins (MBCD), alone (Bru et al. 2006) or combined with the phytohormone methyl jasmonate (MeJA), are used as elicitors (Lijavetzky et al., 2008; Martínez-Esteso et al., 2009, 2011; Belchí-Navarro et al., 2012, 2013; Almagro et al., 2014) due to the combination of the strong induction of biosynthetic genes of shikimate, phenylpropanoid and stilbenoid pathways (Lijavetzky et al. 2008; Almagro et al. 2014) and the ability of cyclodextrins to form inclusion complexes with *t*-R in the extracellular medium (Morales et al. 1998) that protect it from degradation (Jeandet et al. 2021).

Besides biosynthetic genes, a tau-class GST (XM-002275302.2 PREDICTED: Vitis vinifera glutathione S-transferase U10-like; VvGSTU10) was found co-expressed with STS (Martinez-Márquez et al. 2017). GSTs are required for the trafficking and vacuolar accumulation of anthocyanins (Marrs et al., 1995; Alfenito et al., 1998; Kitamura et al., 2004; Li et al., 2011; Sun et al., 2012; Conn et al., 2008; Gomez et al., 2009, 2011), most likely acting as a carrier or “ligandin” rather than a GSH-conjugating enzyme (Mueller and Walbot, 2001). Functional analysis of VvGSTU10 was shown to promote *t*-R transport to the extracellular medium in stably overexpressing grapevine cells (Martinez-Márquez et al. 2017) being the first protein discovered involved in this role.

Different types of ATP binging cassette (ABC) proteins including multidrug resistance (MDR), multidrug resistance associated protein (MRP) and pleiotropic drug resistance (PDR), as well as H^+^-gradient energized multidrug and toxic compound extrusion (MATE) transporters carry out secondary metabolites transport (reviewed in Grotewold et al., 2004; Yazaki et al., 2005; Lefèvre and Boutry 2018; Rahuja et al. 2021). For instance, the alkaloids berberine, catharantine and vincamine (Shitan et al., 2003, 2013; Yu and De Luca 2013; Demessie et al., 2017); the terpenes sclareol, cembrene and caryophyllene (Jasinki et al., 2001; Crouzet et al.,2013, Fu et a., 2017) and the phenolics anthocyanins, glucosylated anthocyanidin, p-coumaryl alcohol, liquiritigenein, 4-coumarate (Gómez et al. 2011; Alejandro et al., 2012; Francisco et al., 2013; Banasiak et al., 2013; Biala et al., 2017; Behrens et al. 2019) are among the secondary metabolites whose transporter proteins have been characterized.

The transcriptomic analysis of MBCD and MeJA elicited grapevine cells displayed a number of genes encoding proteins for transport across membranes of ABC- and MATE-type up regulated and co-expressing with STSs (Almagro et al. 2014), however, to date no one stilbene membrane transporter has been characterized. Here, with the aim of investigating potential transporters candidates involved in the transport of *t*-R accross membranes, we carried out a label-free proteomics analysis of plasma membrane and tonoplast fractions of 72h-elicited grapevine cell cultures. One candidate, VvABCB15 (VIT_214s0066g02320|abc transporter b family member; LOC100854950 –ABC transporter B family member 15; VIT_00032578001) was cloned and functionally analyzed. In vitro *t*-R transport was assessed in transformed yeast microsomes. Moreover, the heterologous system of VvSTS-expressing *Silybum marianum* transgenic cell line (Hidalgo et al., 2017) transformed with VvABCB15, showed extracellular *t*-R levels higher than controls. The functionality of the VvABCB15 transporter was analyzed as well in a homologous system by transient expression in *Vitis* cell cultures. Both elicited and non-elicited transiently transformed lines displayed a greater extracellular *t*-R accumulation, and thus provided for the first-time conclusive evidences of the involvement of VvABCB15 in *t*-R transport to the extracellular medium. Confocal microscopy studies of *N. benthamiana* leaf showed that transiently expressed a VvABCB15-YPF fusion protein displays a plasma membrane localization. The physiological and biotechnological relevance of the results is discussed.

## RESULTS

### Purity of grapevine cells subcellular extracts by MRM analyses

One of the main objectives of this study is to discover candidate transporters involved in the mobilization of *t*-R and likely other monomeric stilbenes, which could be located either in the plasma membrane or tonoplast. For this reason, obtaining fractions enriched in these organelles and as pure as possible is a critical step. For purity check of protein extracts and fractions, westerns or immunoblotting techniques are traditionally used using a set of specific antibodies against organelle-specific proteins. Unfortunately, antibodies for plant research have been developed for proteins of a few model plants thus their utility in the vast majority plant species relying in cross-reactivity is quite limited and scarcely validated. Alternative methods based in targeted proteomics i.e. MRM, have been developed for *Arabidopsis* organelle marker proteins (Parsons and Heazlewood, 2015; Hooper et al. 2017) that surpasses western blotting in multiplexation of the analysis with similar sensitivity. Thus we applied a specific MRM method based on the previously developed SRM markers for *Arabidopsis* organelles suitably adapted to grapevine protein homologs to simultaneously detect and estimate relative organelle abundance, specifically: vacuole, plasma membrane, cytosol and nucleus. Target proteins were: glyceraldehyde-3-phosphate dehydrogenase, monosaccharide transporter tonoplastic, nucleolin like 1, aquaporin PIP1-3-like whose location is cytosol, tonoplast, nucleus and plasma membrane, respectively. Initially, specific tryptic peptides were selected for a unique protein of known localization (see information supporting, table S1). The method was tested on six subcellular fractions enriched by sucrose density gradient centrifugation of a grapevine microsomal fraction (Figure 1A) and two additional samples, crude extract (Ec) and soluble fraction (Fs).

**Figure 1:**
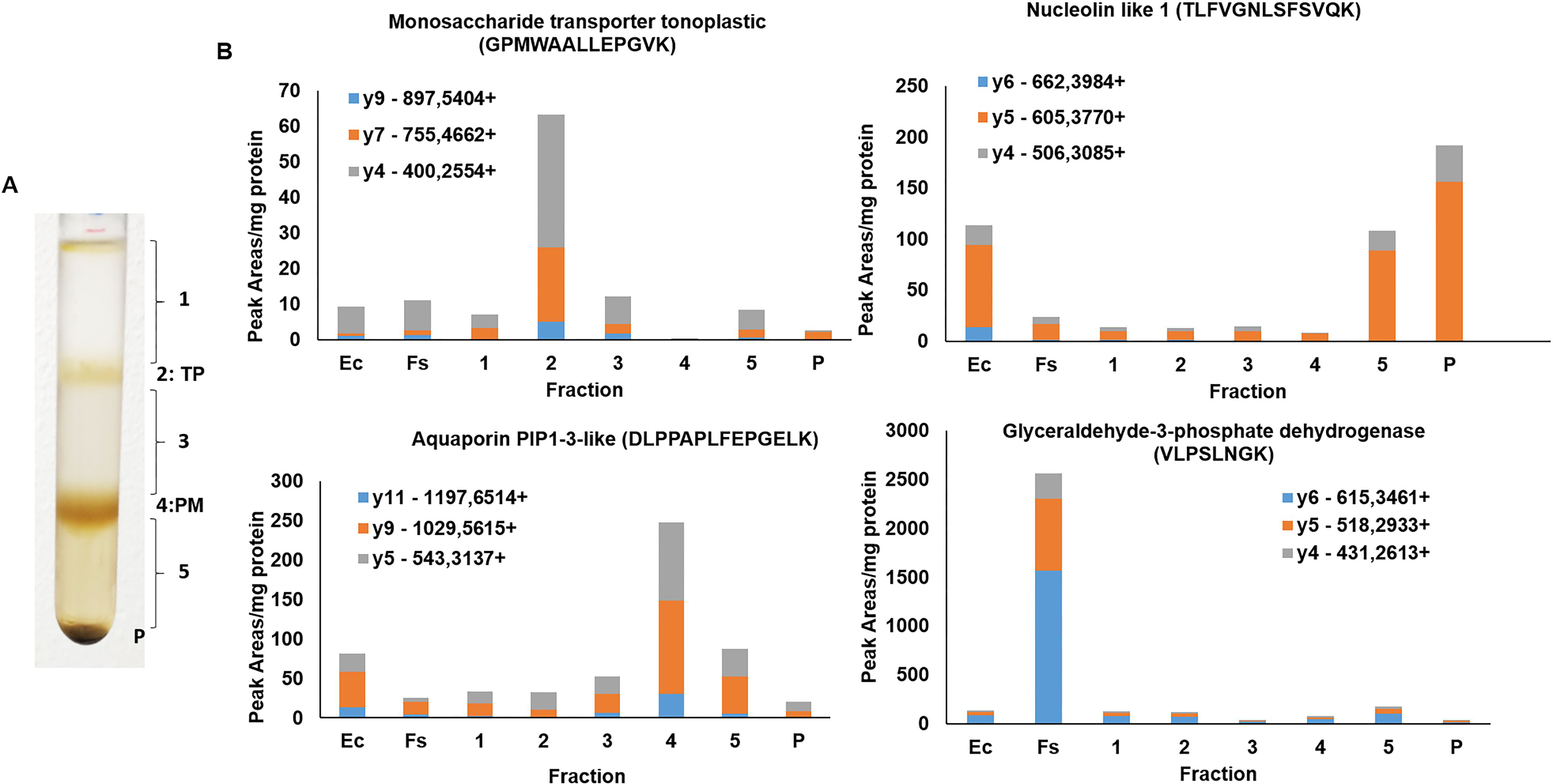
MRM assay in the subcellular fractions enriched by centrifugation density of grapevine microsomal fraction. A) Gradient by centrifugation density of grapevine microsomal fraction. B) Peaks areas of SRM transitions assay normalized with total protein concentration (mg protein) for the different fractions analyzed. Ec, crude extract; Fs, soluble fraction; TP, tonoplast; PM, plasma membrane.

Figure 1B shows peaks areas of different SRM transitions of study normalized by total protein content of the fraction for the different fractions analyzed. According to the results, glyceraldehyde-3-phosphate dehydrogenase is more abundant in the soluble fraction as expected, monosaccharide transporter tonoplastic is enriched in fraction 2 of the gradient, aquaporin PIP1-3-like in fraction 4 and nucleolin like 1 in the crude extract and the heaviest fraction of the gradient and the pellet, as expected. These results confirm that the fraction enriched in tonoplast and plasma membrane are 2 and 4, respectively and, although there is some background of plasma membrane in tonoplast, it represents barely a 10% of the signal in fraction 4. Thus, according to results of figure 1B, it is likely to detect some abundant plasma membrane protein in tonoplast fraction whilst it would be quite unlikely the opposite, or to find nuclear or cytosolic proteins in the tonoplast or plasma membrane fractions. So, the application of the MRM method was successful in checking the purity of the enriched fractions, being effective as well the method for obtaining the enriched fractions of interest.

### Identification of candidate *t*-R transporters

The grapevine cell cultures respond to MBCD and MeJA elicitor treatments with the continuous accumulation of extracellular *t*-R, reaching 3 g/L of culture and above, mainly as the trans-isomer, as described in previous studies (Lijavetzky et al., 2008; Martínez-Esteso et al., 2011a, Martinez-Marquez et al., 2017). Quantitative proteomic analysis of whole cell extracts has revealed the up-regulation of t-R biosynthetic pathway enzymes PAL and STS as well as unanticipated elicitor-responding proteins such as tau class GST (Martínez-Esteso et al., 2011b, Martinez-Marquez et al., 2017). The functional analysis of the latter provided evidences of the first protein involved in transport of *t*-R to the extracellular medium (Martinez-Marquez et al., 2017). To broadly explore the expression profiles of proteins potentially involved in the *t*-R transport, a label-free proteomic experiment of enriched extracts in tonoplast or plasma membrane of 72 h elicited grapevine cell cultures were carried out (Figure 2).

**Figure 2.**
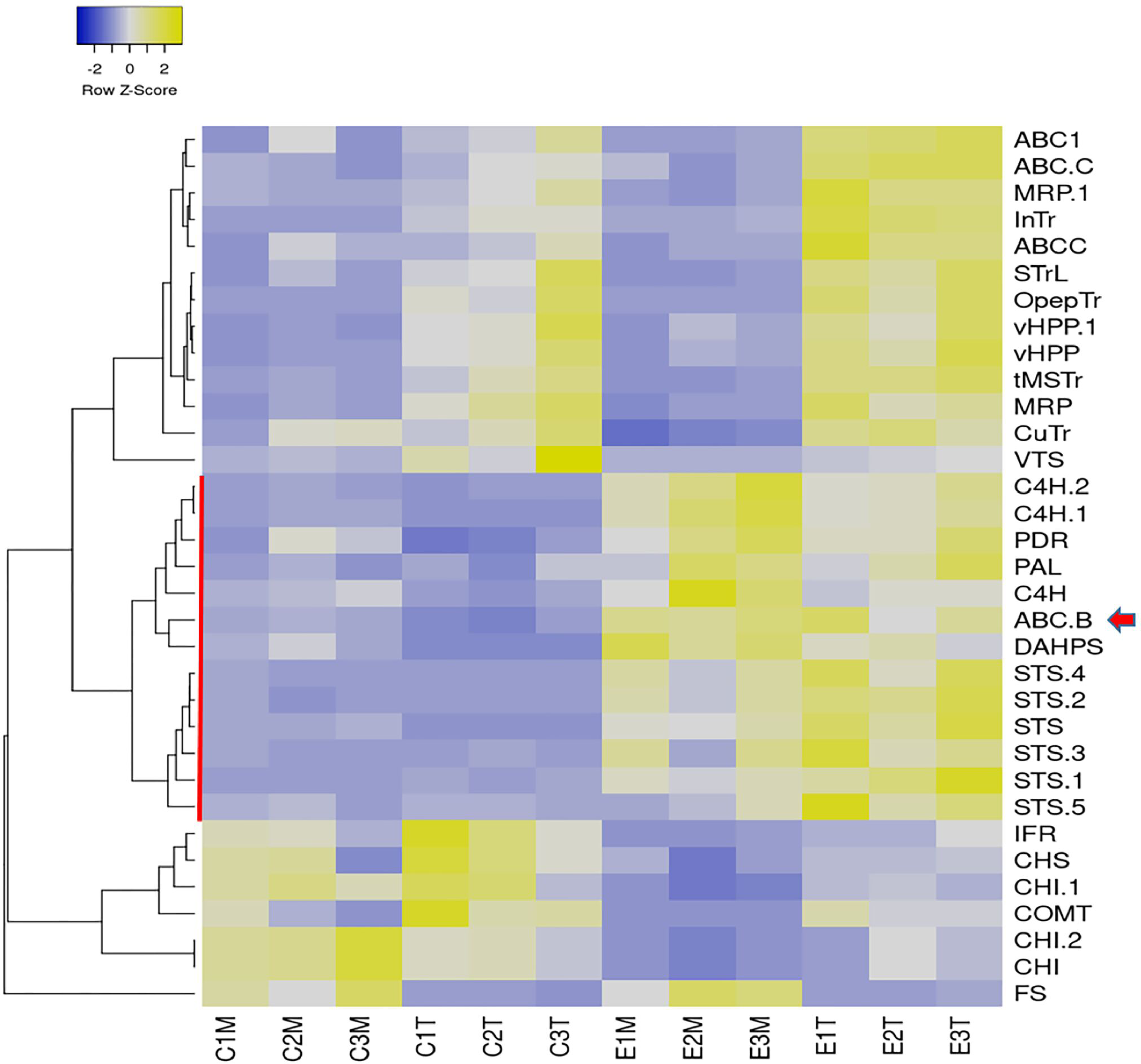
Label-free proteomic analysis of control (C#) elicited (E#) plasma membrane (M) and tonoplast (T) fractions. Normalized abundance heatmap of selected proteins filtered by ANOVA p<0,02 and fold change>4, involved in biosynthesis of stilbenes, flavonoids and lignans and transport across membranes: 3-deoxy-D-arabino-heptulosonate 7-phosphate synthase (DAHPS), phenylalanine ammonia-lyase (PAL), cinnamate-4-hydroxylase (C4H), stilbene synthase (STS), chalcone synthase (CHS), chalcone isomerase (CHI), flavonol synthase (FS), isoflavone reductase homolog (IFR), caffeic acid O-methyltransferase (COMT), vacuolar H+-translocating inorganic pyrophosphatase (vHPP), tonoplastic monosaccharide transporter (tMSTr), ABC transporter B family member 15 (ABC B: VvABCB15), ABC transporter C family member 3-like and family member 4-like (ABC C), ABC1 family protein (ABC1), pleiotropic drug resistance protein 1-like (PDR), multidrug resistance protein MATE (MRP), sugar transporter erd6-like 6 (STrL), oligopeptide transporter 7-like (OpepTr), vesicle transport V-snare 13 (VTS), inositol transporter (InTr), copper transporter (CuTr). Figure prepared with Heatmapper (Babicki et al. 2016) using average linkage for clustering and Pearson as distance measurement method.

In total, 1637 proteins were identified of which 146 were found with significant differential abundance under the elicitation conditions (ANOVA p <0.02 and fold change> 4).

Figure 1-suppl. summarizes the label-free proteomic analysis of control and MBCD/ MeJA-elicited plasma membrane (PM) and tonoplast (TP) fractions. Hierarchical clustering analysis of the abundance patterns distance (Fig. 1A-suppl) was set to classify the profiles into six color-coded groups (Fig. 1B-suppl). The same color code was used for the PCA bi-plot (Fig. 1C-suppl). The up-regulated in elicitation treatments vs control group (green) is the one that likely contains candidates for *t*-R transport across membranes. This green target group contained proteins involved in the biosynthesis of stilbenes (DAHPS, PAL, C4H, STS), an ABC B class transporter and a pleiotropic drug resistance protein belonging to the ABC G class transporter. Proteins that are competitors of STS for metabolic precursors (CHS, CHI, COMT, IFR) belonged to the down-regulated in elicitation treatments group (pink) or (FS) more abundant in PM-enriched (orange). In order to better select *t*-R transport candidates, we focused on proteins involved in biosynthesis of stilbenes, flavonoids and lignans and transport out of the 146 deregulated group. As seen in Figure 2, membrane transporters including known tonoplast proteins (vHPP, tMSTr), characterized (InTr, CuTr, OpepTr, STrL) and uncharacterized transporters (ABC C, ABC1, MRP-MATE) clustered as up-regulated in tonoplast fraction with or without elicitation, while the ABC B and PDR transporters clustered with stilbene biosynthesis enzymes, up-regulated in elicited groups.

According to these results, the ABC B and PDR are a good candidates for the mobilization of *t*-R towards the extracellular medium in response to elicitors. In order to try to discriminate between both we studied the correlation between the fold-change at the level of protein (data obtained here) with that of transcripts, reported after 24h of elicitation with MeJA + MBCD in *V. vinifera* cv Monastrell (Almagro et al 2014). For comparison, we also included the GST U10-class involved in *t*-R extracellular accumulation (Martinez-Marquez et al. 2017). To cancel the effect of differential expression due to fractionation the comparison was done for each fraction, PM and TP, separately. As shown in Figure 3, ABC B, as well as stilbene biosynthesis products (DAHPS, PAL, C4H, STS) correlated positively, however PDR transcripts encoding that PDR protein were not found in Almagro’s et al. 2014 report. Other transporters showed a poor correlation, likely due to the fraction cancellation effect.

**Figure 3.**
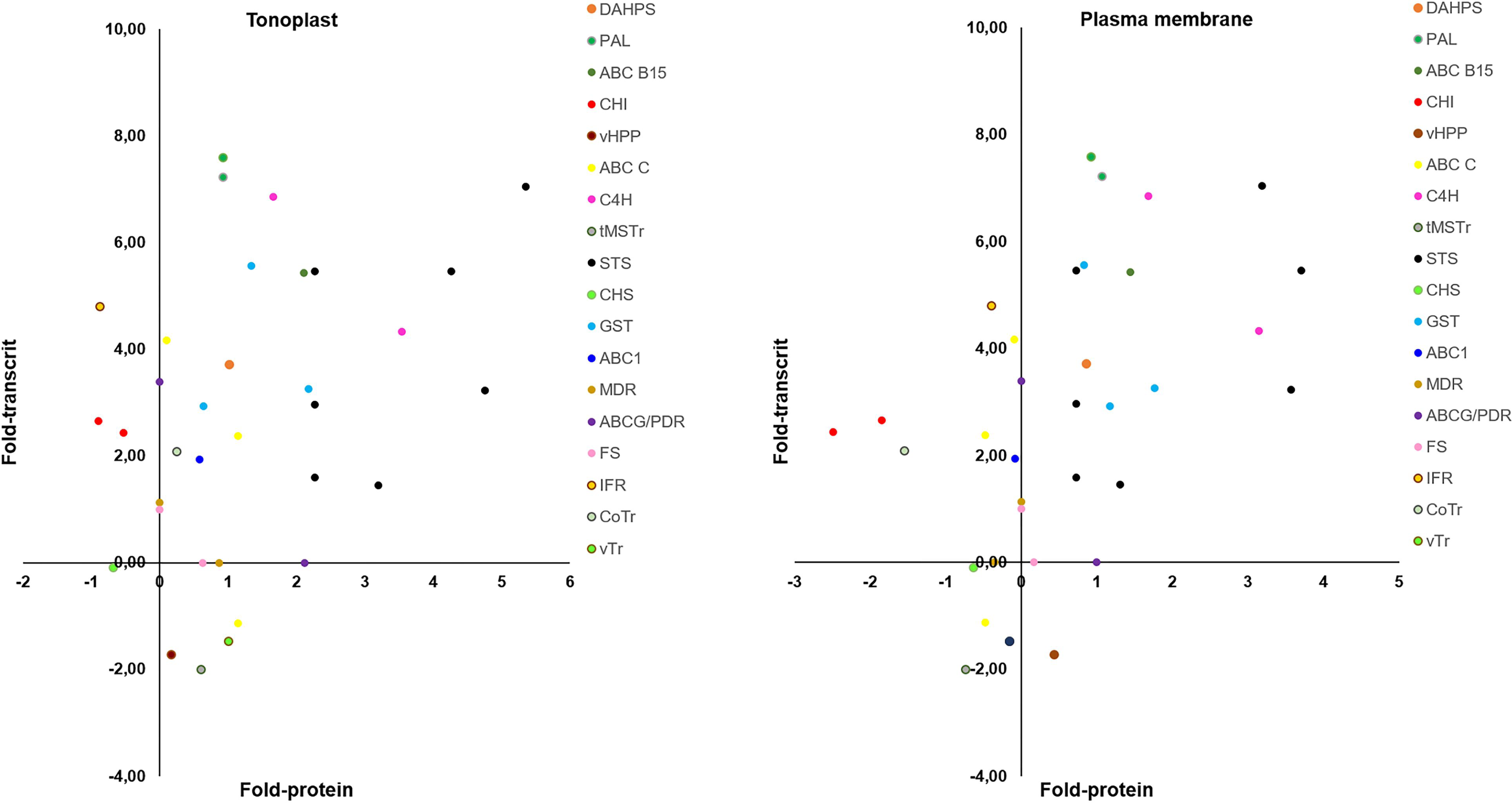
Comparison of transcript and protein changes in response to elicitation treatment with 50mM MBCD and 100µM. Transcript and protein fold-change presented as the log2 of the ratio of elicited:control conditions.Transcript fold change was obtained by microarrays in Vitis Vinifera cv. Monastrell at 24h of elicitation treatment (p<0,05 and fold change>2; see Supplementary information Almagro et al., 2015). Protein fold change was obtained by label-free proteomic analysis in tonoplast (A) plasma membrane (B) fractions of Vitis Vinifera cv. Gamay at 72h of elicitation treatment (p<0,02 and fold change>4; see Supplementary information). Fold-change for biosynthesis of stilbenes, flavonoids and lignans and transport across membranes. 3-deoxy-D-arabino-heptulosonate 7-phosphate synthase (DAHPS), phenylalanine ammonia-lyase (PAL), cinnamate-4-hydroxylase (C4H), stilbene synthase (STS), glutation-S-transferase (GST), chalcone synthase (CHS), chalcone isomerase (CHI), flavonol synthase (FS), isoflavone reductase homolog (IFR), vacuolar H+-translocating inorganic pyrophosphatase (vHPP), tonoplastic monosaccharide transporter (tMSTr), ABC transporter B family member 15 (ABC B: VvABCB15), ABC transporter C family member 3-like and family member 4-like (ABC C), ABC1 family protein (ABC1), pleiotropic drug resistance protein 1-like (PDR), multidrug resistance protein MATE (MRP),vesicle transport V-snare 13 (vTr), copper transporter (CoTr). Fold changes correspondent to transport across membranes are marked with a circle.

Taking into account the above results we selected the ABC B (VvABCB15) transporter for functional characterization.

### VvABCB15 localizes to the plasma membrane

To determine the subcellular localization of VvABCB15, the full-length gene was C-terminally fused to yellow fluorescent protein (YFP) and expressed under the control of the CAMV 35S promoter (P35S:VvABCB15-YFP). A plasma membrane aquaporin fusion PIP2A-CFP (for cyan fluorescent protein) was used as a control for plasma membrane localization (Nelson et al., 2007).

Four days agroinfiltrated *N. benthamiana* leaves observed under confocal microscopy showed the YFP signal as a band at the cell periphery, indicating that VvABCB15 could be located in the plasma membrane. As shown, the distribution of fluorescence for both PIP2A-CFP (Fig 4A,D) and VvABCB15-YFP (Fig 4B,E) are found at the periphery of the cells when a single optical section is observed. Furthermore, merged images show that the two individual signals coincide almost exactly (Fig 4C,F) consistent with a plasma membrane co-localization (0,68 pearson coefficient). Results provide clear evidence for plasma membrane localization of VvABCB15.

**Figure 4:**
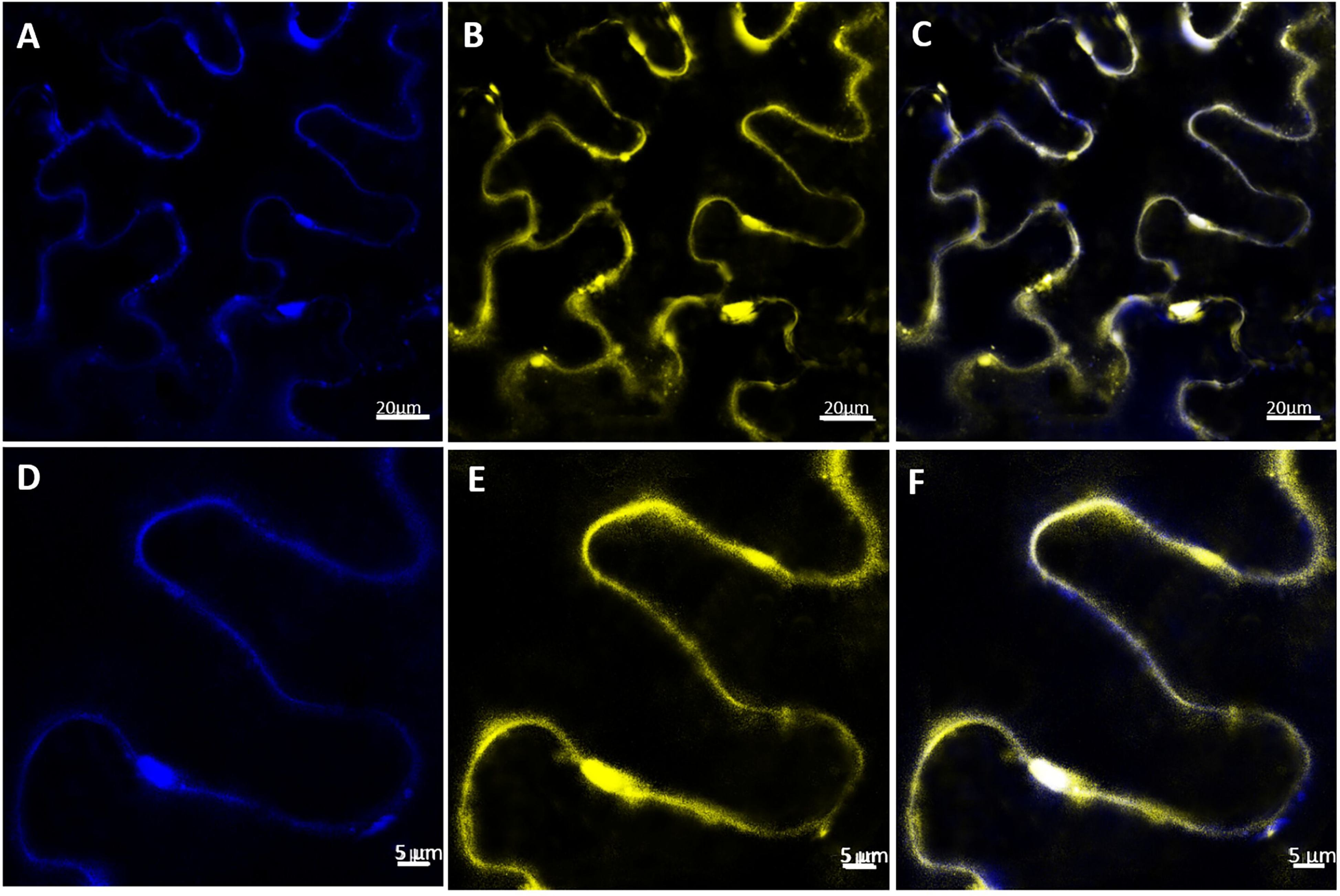
Subcellular localization of VvABCB15 transporter in *N. benthamiana* leaves by confocal microscopy four days after agroinfiltration. (A, D) PIP2-CFP expression in *N. benthamiana* leaves (B, E) VvABCB15-YFP expression in N. benthamiana leaves (C,F) merged images of one optical section.

### Functional characterization of VvABCB15 as a *t*-R transporter in yeast microsomes

It is known that ABC transporters can transport a large number of chemically unrelated compounds (Kang et al., 2011). Following the observation that VvABCB15 co-expressed with *t*-R biosynthetic proteins in response to elicitors and that it is localized to the plasma membrane, we undertook the evaluation of its functionality as stilbene transporter in both heterologous and homologous systems as an ultimate prove of its biological role. For this purpose, we ordered a synthetic version the VvABCB15 gene cloned into the pESC-URA-cMyc plasmid for expression in *S. cerevisiae* fused in C-terminal to a Myc tag. The presence of VvABCB15 in transformed yeast microsomal membrane vesicles was confirmed by western blot with antiMyc antibody (Fig. 5A). *t*-R is the most abundant stilbene in extracellular grapevine cell cultures but also the cis isomer and dimers known as viniferins may accumulate to a significant extent (Lijavetzky et al., 2008; Martínez-Esteso et al., 2011a, Martinez-Marquez et al., 2017). Thus, transport assays in the presence of *t*-R and MBCD as a *t*-R carrier were conducted using the rapid filtration technique as reported by Tommasini et al. (1996). Targeted quantitative analysis of stilbenoids by MRM was used to quantify the stilbenes taken up into the vesicles.

**Figure 5:**
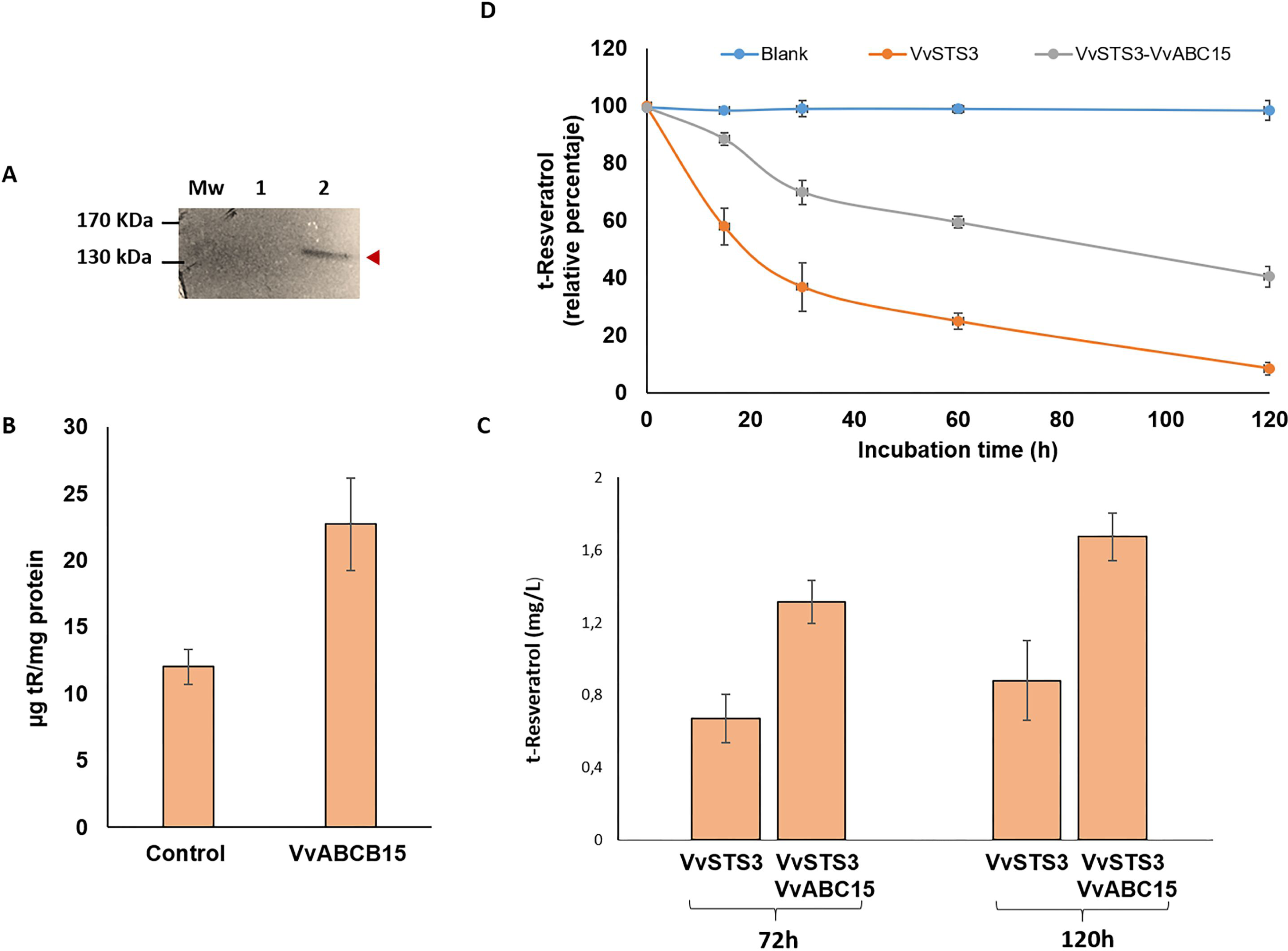
Functional analyses in heterologous systems. A) Expression of the VvABCB15 recombinant protein in yeast cells. Expression of the Myc-tag fusion proteins was confirmed by a Western blot analysis with an anti-Myc-tag antibody. B) Net t-R uptake into microsomes isolated from yeast cells transformed either with the empty vector (control) or with VvABCB15 after 30 min incubated with 2.98mM of t-R. Results are presented as mean values ± SE of two independent uptake experiments. C) Extracellular accumulation of t-R in the stable transgenic silybum cell suspensions in the presence of 5mM elicitor MBCD for 72h and 120 h. Data are the mean of three independent replicates ± SD. D) Residual level of exogenously added t-R to STS-(orange) and STS+ABCB15-(grey) expressing *Sylibum marianum* transgenic cell lines. As control, residual level of t-R added to the medium but without cells (blue).

In all the control assays, carried out with yeast microsomes that do not express ABC transporter, we observed an amount of *t*-R possibly due to either passive or nonspecific endogenous transport processes. However, significantly higher uptake of *t*-R into yeast microsomes expressing VvABCB15 was observed (Fig. 5B). Also controls without microsomes showed that a measurable amount of *t*-R bounded to the filter that was substracted in both VvABCB15- and non VvABCB15-expressing microsome assays. These observations in a heterologous system are the first proof of the functionality of the VvABCB15 transporter in the mobilization of *t*-R.

### Functional analyses by heterologous stable expression in VvSTS3-expressing *S. marianum* cell culture

VvSTS3-expressing *S. marianum* transgenic cell line represent a heterologous system for *t*-R production under conditions of elicitation with MBCD (Hidalgo et al., 2017). For this reason, and with the objective of evaluating the functionality of the VvABCB15 transporter in a plant heterologous system, this transgenic cell line was transformed with the construction harbouring a TU for VvABCB15 (Figure 4D).

Control, STS3-expressing, and test, doubly expressing VvSTS3+VvABCB15, *S. marianum* cell suspensions were incubated in the absence and presence of 5mM MBCD. As seen in the chromatograms of the extracellular media, *t*-R and an uncharacterized viniferin dimer can be detected in non MBCD treated suspensions as tiny peaks in control (Figure 2-suppl.A) but as prominent peaks in STS3+VvABCB15 (Figure 2-suppl.B). On the other hand, MBCD treatment strongly promotes the accumulation of these compounds in the extracellular medium, as described by Hidalgo et al. (2017), but the *t*-R and viniferin peaks are also larger in STS3+ABCB15 than in control (Figure 2-suppl.C, D). Figure 5C shows the total accumulation of extracellular *t*-R after 72 and 120 hours in the presence of 5 mM MBCD, in VvSTS3+VvABCB15 transformed line and VvSTS3 control line. As expected from the quantitative analysis of chromatograms, accumulation in the VvSTS3+VvABCB15 transformed cell suspensions was about 1.9-fold higher in each of the MBCD treatments than in control.

Likewise, it has been reported that *t*-R added to a grapevine cell suspension disappear after some hours (Morales et al. 1998) due to cell uptake and metabolism. Here, *t*-R was added externally to *S. marianum* cells expressing VvSTS3 or VvSTS3+VvABCB15 to follow its evolution with time under the hypothesis that an active outwards transport by VvABCB15 would secrete the *t-*R took up by cells thus keeping its extracellular level higher than in control STS3-expressing cells. As seen in Figure 5D, that hypothesis was confirmed. Taken altogether, these results obtained in the *S. marianum* heterologous system provides further evidence of the functionality of VvABCB15 in the transport of free *t*-R.

### Functional analyses by homologous transient expression in *Vitis* cell culture

The increased extracellular accumulation of *t*-R under non elicited conditions of *Vitis vinifera* cv. Gamay cells was used as a functional assay for putative candidate genes/proteins involved in *t*-R transport out of the cell (Martinez-Marquez et al., 2017). Gamay cells are able to constitutively synthesize stilbenes (*c*-Piceid and *t*-Piceid, and a modest *t*-R) and store them within cells during normal growth conditions (Martínez-Esteso et al., 2011, Martinez-Marquez et al., 2017). The occurrence of an active stilbene transport system would lead to the presence of the stilbene compounds outside of the cells at higher level, as it was shown for the stable overexpression of VvGSTU10 (Martinez-Marquez et al. 2017). Here, transient expression experiments were carried out in Gamay grapevine cell cultures in the presence of adsorbent compounds PVP or βCD without elicitor effect (Bru et al., 2006; Martinez-Marquez et al., 2017), to stabilize the cell-secreted *t*-R. Five days after agroinfection of the cell culture, extracellular *t*-R content was analysed (Figure 7A). Small amounts of *t*-R, below 5 mg/L, were detected in the extracellular medium of the wild cells and in the transformation control GFP-expressing cells due to both the basal production of stilbenes in Gamay cells and the stabilizing effect of both PVP or BCD (Martinez-Marquez et al., 2017). In the positive transport control VvGSTU10-expressing cells *t*-R extracellular levels reached above 15 mg/L. The transient expression of VvABCB15 resulted in *t*-R extracellular levels above 20 mg/L and the joint transient expression of both VvGSTU10+VvABCB15 lead to even higher levels, between 25 and 30 mg/L. This result demonstrates that under non-elicitation conditions, *t*-R transport towards the extracellular medium occurred to a much greater extent in VvABCB15 transiently transformed than in control or in wild cells, and that co-expression of both VvGSTU102 and VvABCB15 increases the transport capacity of the grapevine cells as compared to their individual counterparts. Results are virtually similar irrespective of the adsorbent compound used. This result is highly consistent with both its localization in plasma membrane and the *t*-R mobilization activity demonstrated in yeast microsomes.

It is well known that the use of MBCD as an elicitor, either alone or combined with MeJA, usually at 50mM and 100µM respectively, increases strongly the expression of genes of stilbene biosynthetic pathway and genes involved in *t*-R transport such as VvGSTU10 (Almagro et al., 2013, Martinez-Marquez et al., 2017), leading to a continuous increase in extracellular *t*-R and steady-state levels in the intracellular compartments. Under mild elicitation conditions (5mM MBCD) the effect of stable overexpression of VvGSTU10 on t-R extracellular accumulation could be clearly seen above the background effect of the elicitor (Martinez-Marquez et al., 2017). Thus we also analysed the VvGSTU10 and VvABCB15 transient expression in Gamay cells under mild elicitation. Figure 7B shows the amount of extracellular *t*-R accumulated after 72h of incubation with 5mM of MBCD. As expected from the elicitor activity of MBCD, abundant extracellular *t*-R was found in all cell suspensions tested, but accumulation in the wild type and GFP-expressing transiently transgenic cell suspensions were much lower than in the VvGSTU10, VvABCB15 or VvGSTU10+VvABCB15 transiently transformed cell suspensions. Differences in the accumulated *t*-R between transformation control GFP-expressing and wild-type were not significant, and reached between 50 and 70 mg/L. Separate expression of either VvGSTU10 or VvABCB15 led to higher levels, between 130 and 140 mg/L but without significant differences between them. Interestingly, their co-expression causes a further increase, reaching above 200 mg/L. This result is quite consistent with that obtained above under non-elicitation conditions and highlights the finding that both VvGSTU10 and VvABCB15 cooperate in the transport and extracellular accumulation of *t*-R in grapevine cells.

On the other hand, several studies have shown an increase in the accumulation of *t*-R upon the heterologous expression or overexpression of STS genes in different plant systems (Hain 1990, 1993; Donnez et al., 2009; Kiselev and Dubrovina et al., 2021) occurring in the extracellular medium in case of cell or tissue cultures (Hidalgo et al., 2017a, b; Chu et al., 2017). In this sense, quantitative changes in intra- and extra-cellular *t*-R in STS-overexpressing cells associated to co-expression of a candidate can also be used as functional assay for *t*-R transport. Here we have overexpressed VvSTS1 and VvABCB15 under control of the constitutive promoter p35S in grapevine cells through transient transformation and quantified stilbenes inside and outside the cells by MRM in a multiplexed analysis, including *t*-R, *c*-R and their glycosylated forms, i.e., piceid (*t*-Pc). In addition, VvSTS3 expression has also been handled using the light-switchable promoter pHyH (Gangappa and Botto, 2016) to better dissociate the effects of infection from those of VvSTS1 overexpression on stilbene accumulation (see Fig 6 for constructs). Assays include two controls, namely the wild type culture, and the transient expression of GFP under pRolD promoter as a negative control of the agroinfection. Figure 8 shows the amount of the extracellular (Fig 8A) and intracellular (Fig 8B) stilbenoids accumulated after 5 days of infection either in darkness or under photoperiod. As mentioned above, wild-type Gamay cells produce constitutively mainly the glycosylated form piceid and a little of the free *t*-R, all intracellular. The transformation control expressing GFP shows a significant increase as compared to wild-type, particularly of the free form in both the cells and the extracellular medium. Either wild-type or GFP-expressing cells do not show differences between darkness and photoperiod conditions. In darkness, the stilbene accumulation in pHyH-VvSTS3 transformed cells is slightly higher than in the transformation negative control. The above results indicate that the light regime has not a background effect on stilbene production, and that transformation itself causes a significant increase in the basal levels of free stilbenes, and a little effect on the glycosylated forms. In pHyH-VvSTS3 transformed cells, photoperiod conditions lead to a strong increase in stilbene accumulation as compared to darkness due to VvSTS3 expression. This result suggests that there must be constitutive transporters that facilitate the leave of the readily VvSTS3-produced *t*-R. The co-transformation pHyH-VvSTS3/p35S-VvABCB15 compared to pHyH-VvSTS3 causes an increase in extracellular *t*-R in both conditions (1.64-fold for darkness and 2.1-fold photoperiod) and simultaneously, a decrease in intracellular stilbenes in both conditions, darkness (0.93-fold for *t*-R and 0.83-fold piceid) and photoperiod (0.77-fold for *t*-R and 0.64-fold piceid). The same phenomenon is observed when the expression of STS is constitutive, that is, co-transformation p35S-VvSTS3/p35S-VvABCB15 compared to p35S-VvSTS3 causes an increase in extracellular *t*-R of 1.74-fold and simultaneously, a decrease in intracellular stilbenes (0.71-fold for *t*-R and 0.69-fold piceid). These results can be interpreted as the effect of VvABCB15 overexpression that increases the rate of outward transport of the readily VvSTS3-produced free *t*-R, thus competing with the glycosylation reaction (supposedly unchanged) that keeps the *t*-R inside the cells as piceid. Consequently, the steady state level of *t*-R and the accumulated piceid decrease within cells whilst extracellular levels of free stilbenes increase.

**Figure 6:**
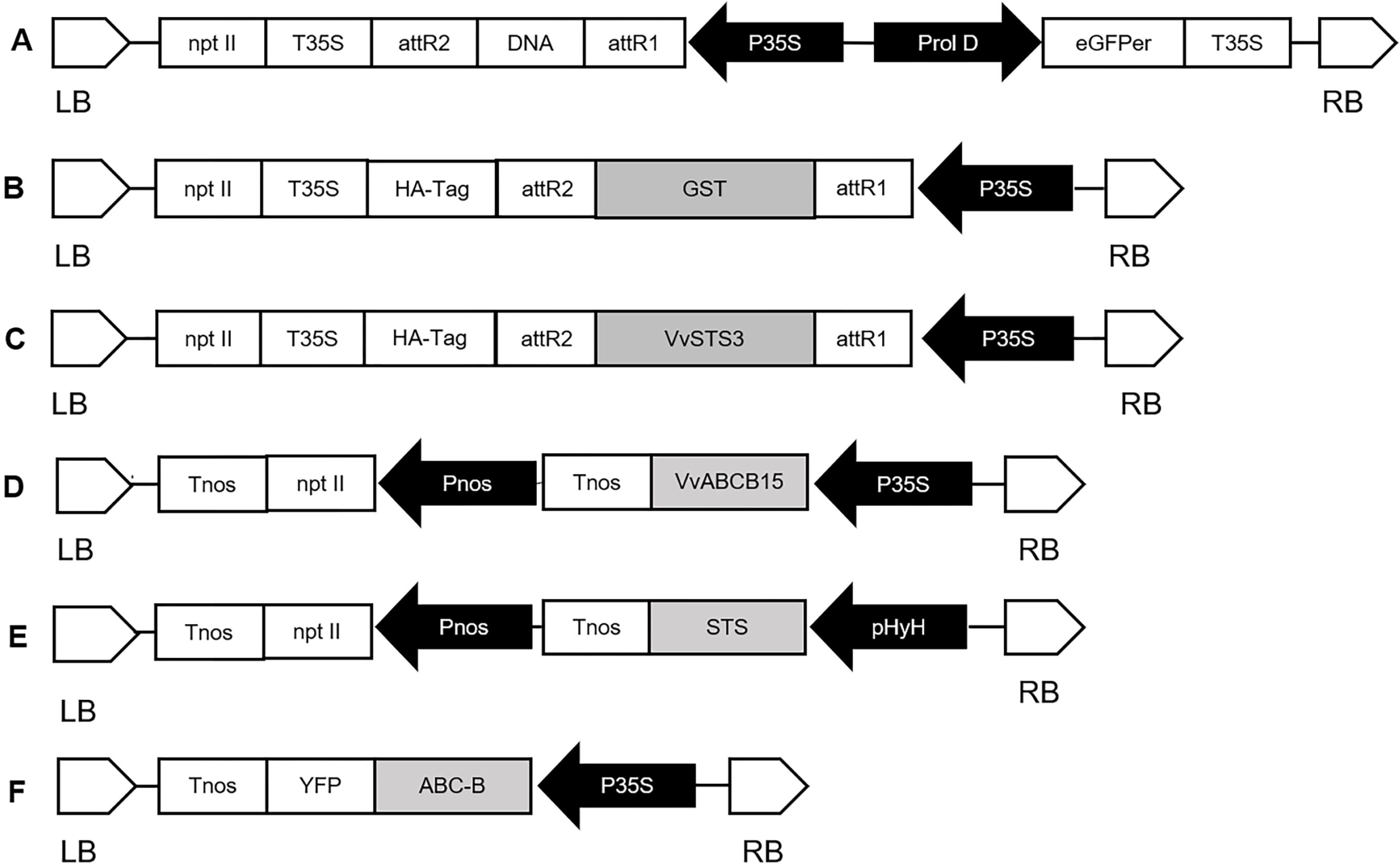
Schematic diagram of the expression constructs used for transient transformation experiments. P35S, cauliflower mosaic virus (CaMV) 35S promoter; Pnos, nopaline synthase promoter; pHyH, HyH promoter; nptII, neomycin phosphotransferase; VvSTS3, Stilbene synthase; VvGSTU10, Glutathione-S-transferase; VVABCB15, ATP binding cassette B 15; eGFPer, Green fluorescent protein with ER signal peptide sequence; YFP, Yellow fluorescent protein, T35S, CaMV 35S terminator; Tnos, nopaline synthase terminator; LB, left border; RB, right border. (A) pK7WG2D-GFP (Martinez-Marquez et al., 2015); (B) pJCV52-VvGSTU10 (Martinez-Marquez et al., 2017); (C) pJCV52-VvSTS (Hidalgo et al., 2017); (D) pDGB-VvABCB15; (E) pDGB-VvSTS; (F) pDGB-VvABCB15-YFP.

**Figure 7:**
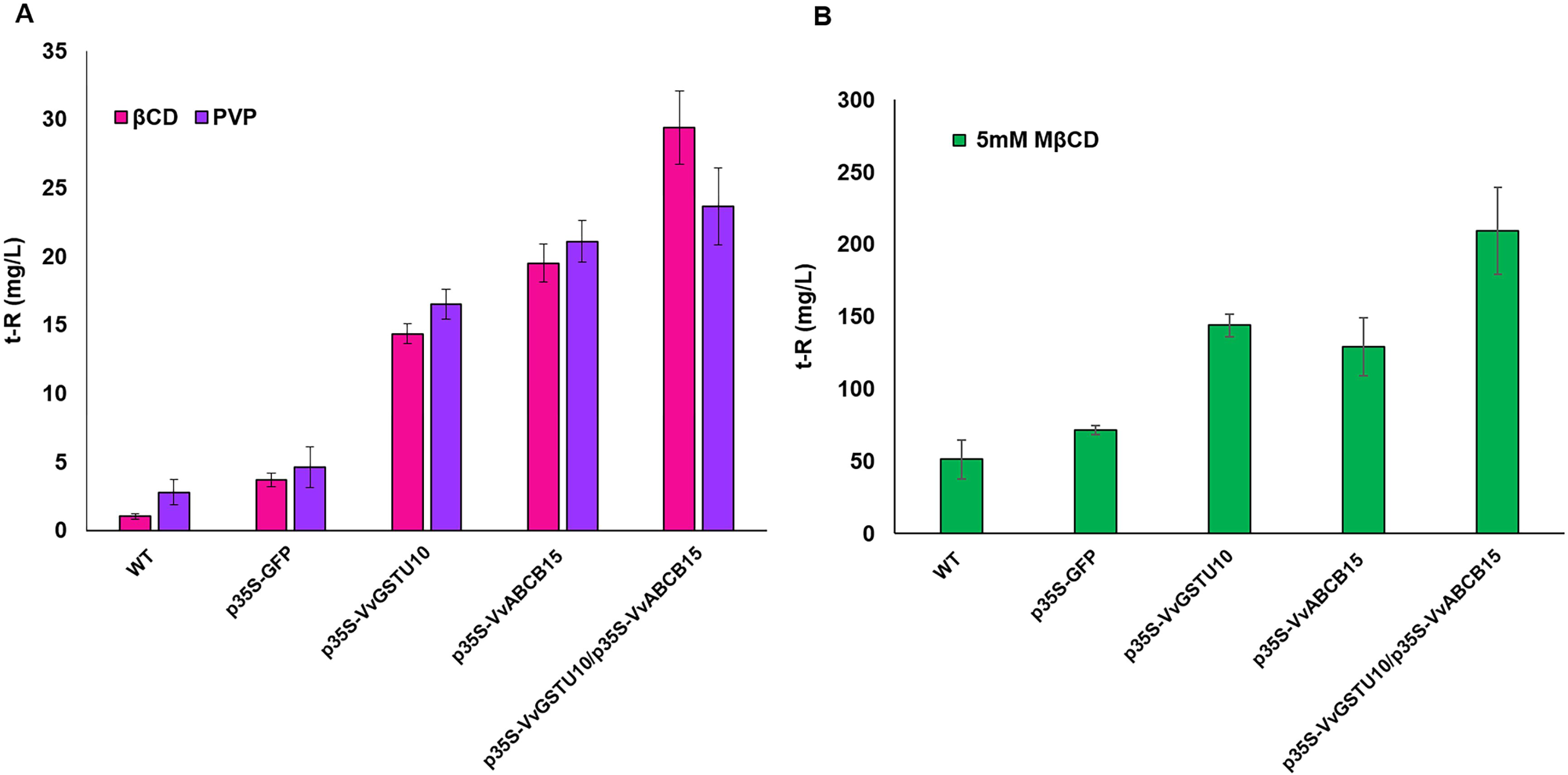
Extracellular accumulation of t-R in grapevine cell suspensions transiently transformed in the presence of PVP or BCD (A) and elicitor MBCD (B). Effect of absorbent compound PVP (blue) and BCD (pink) on the extracellular t-R accumulation at 5 days after agroinfectation. Effect of 5mM elicitor MBCD (green) on the extracellular t-R accumulation for 72h after agroinfectation. Data are the mean of three independent replicates ± SD.

**Figure 8:**
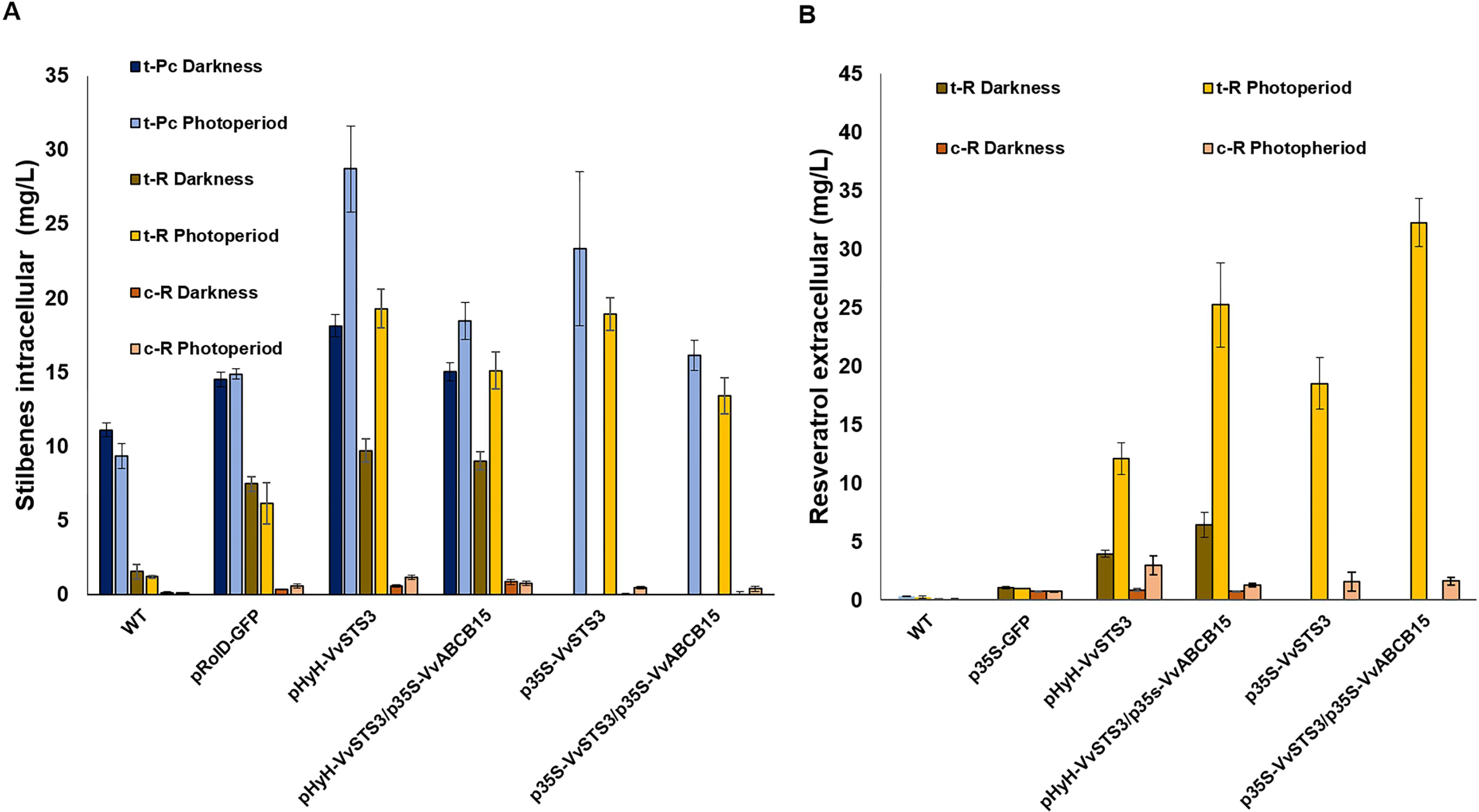
Stilbenes accumulation 5 days after agroinfection in grapevine cell suspensions transiently transformed. A) Effect on the extracellular of trans-resveratrol (t-R) and cis-resveratrol (c-R) in darkness and photoperiod. B) Effect on the intracellular of t-R, c-R and trans-piceid (t-Pc) in darkness and photoperiod. Data are the mean of three independent replicates ± SD.

*c*-R content did not change compared to the wild-type or GFP-expressing cells, or changed only slightly for pHyH-VvSTS3 transformed cells intracellularly and for all VvSTS3 or VvSTS3+VvABCB15 transformed cells extracellularly, especially in photoperiod

## DISCUSSION

Phenolic secondary metabolites such as stilbenes, play key biological roles in plant defence. These may pre-exist in high content or may be synthesized after microbial attack being part of both constitutive and inducible defence responses respectively (Chong et al., 2009; Plant Science 177 (2009) 143–155). Their translocation to the extracellular compartment from the internal pools is essential as the first line of defence. This highlights the physiological importance of stilbenes transport to be biologically efficient.

Grapevine cell cultures show synthesis and accumulation of *t*-R in the extracellular compartment in response to an array of biotic and abiotic elicitors (Liswidowati et al., 1991; Calderon et al., 1993; Krisa et al., 1999; Belhadj et al., 2008; Yue et al., 2011; Tassoni et al., 2005; Ferri et al., 2009; Morales et al., 1998; Bru et al., 2006; Zamboni et al., 2006; Lijavetzky et al., 2008; Martínez-Esteso et al., 2009, 2011b; Belchí-Navarro et al., 2012, 2013; Almagro et al., 2014; Almagro et al., 2015), being ideal biological platforms for the study of resveratrol transport. In this sense, Martinez-Marquez et al., 2017 demonstrated the involvement of VvGSTU10 in *t*-R transport to the extracellular medium in grapevine cell culture elicited with MBCD combined with MeJA. However, despite their utmost physiological and biotechnological relevance, the transport pathways of *t*-R and other stilbenes to the extracellular medium in grapevine cells are completely unknown. The present work is thus the first study to aim to find membrane transporters involved in this transport process.

The proteomic experiment on plasma membrane and tonoplast fractions of 72h-elicited grapevine cell cultures (Figure 2) has allowed the discovery of VvABCB15 as a candidate involved in this process. Previously, the purity of the tonoplast and membrane-enriched fraction was checked by MRM method (Figure 1). We find several transporters that fit the expression profile of biosynthetic enzymes (Figure 2). However, the comparison of transcript (Almagro et al., 2015) and protein changes in response to elicitation treatment with 50mM MBCD and 100µM MeJA (Figure 3) led us to select an ABC-B type transporter.

Specifically, ATP-binding cassette (ABC) proteins are multidomain transmembrane proteins that use the energy obtained from ATP hydrolysis to translocate molecules like xenobiotics, hormones, sugars, amino acids, ions, primary and secondary metabolites among others, involving in essential phyological processes such as nutrition, development, responses to biotic and abiotic stress, interaction with the environment, and mainly in transmembrane transport (Lefèvre and Bountry 2018; Nogia and Pratap 2021). Generally, ABC transporters possesses two types of domains, namely cytosolic nucleotide-binding domain (NBD)/ATP binding domain and transmembrane domain (TMD) (Sharom et al., 2011). Based on the combination and number of TMDs/NBDs, the ABC transporter family has often been grouped into nine subfamilies viz. ABCA, ABCB, ABCC, ABCD, ABCE, ABCF, ABCG, ABCH, and ABCI (Verrier et al., 2008) amongst which ABCH has not been identified in plants. Their involvement in the trafficking of secondary metabolites has been well demonstrated for alkaloids (Shitan et al., 2003, 2013), terpenes (Jasinki et al., 2001; Yu and De Luca 2013; Demessie et al., 2017; Fu et a., 2017), phenolic compounds as glucosylated anthocyanidin (Francisco et al., 2013; Behrens et al. 2019) or monolignol (Alejandro et al., 2012), and volatile compounds (Adebesin et al., 2017). However, no membrane transporter has been described as a protein involved in the mobilization of stilbenes or more specifically *t*-R.

So far, 120 putative ABC members have been identified in grapevine (*Vitis vinifera*) (Çakır and Kılıçkaya, 2013). The main feature of the Vitis ABC superfamily is the presence of several large subfamilies, among which is included ABCG (pleiotropic drug resistance and white-brown complex homolog, PDR), ABCC (multidrug resistance-associated protein, MRP) and ABCB (multi-drug resistance/P-glycoprotein, MDR/PGP). Plant MRP/ABCCs subfamily has been proposed to be primarily involved in the vacuolar sequestration of potentially toxic metabolites (Klein et al., 2006). In fact, the expression profiles of ABCC subfamily transporters detected in our label-free proteomic experiment were higher in the tonoplast (Figure 2), reason for which they were not selected as candidates for the transport of stilbenes outside the cell. Members of ABCG family are expressed in plants in response to various biotic and abiotic stresses; and play important roles in detoxification processes, preventing water loss, transport of phytohormones, and secondary metabolites (Alav et al., 2021). A PDR belonging to the ABC G class transporter, which fit the expression profiles of *t*-R biosynthetic enzymes (Figure 2), was discovered along with an ABC B class transporter. Because PDR transcripts encoding that PDR protein were not found in Almagro’s et al. 2014 report we select the ABC-B transport as the only candidate involved in *t*-R transport outside of *Vitis* cells. Only a few ABCB transports have been extensively characterized in plants and shown to catalyze the transport of structurally diverse substrates, such as phytohormones, xenobiotics and secondary metabolites (Bailly et al., 2012). In addition, gene expression studies made by Kaneda et al., (2010) show a correlation between expression of a subset of genes encoding ATP-binding cassette (ABC) transporters, especially in the ABCB, and phenylpropanoid biosynthetic genes, that are involved in stilbenes biosynthesis.

The subcellular localization studies carried out herein (Figure 4) showed that VvABCB15 could be a plasma membrane transporter due to colocalization with a membrane protein a plasma membrane aquaporin PIP2A.

The functional characterization VvABCB15 was successfully performed in both heterologous and homologous systems. Wild yeast and *Silybum marianum* cell cultures stably expressing VvSTS3 were used as heterologous systems. In both cases, our results showed that resveratrol transport occurs more intensively in both VvABCB15-transformed heterologous systems than in their controls (Figure 5). It is known that, in addition to the need of ATP for the functioning of ABC transporters, GSH is co-transported with the specific substrates in some cases (König et al., 2003). Thus, for studies with yeast microsomes, we performed the transport assays including both ATP and GSH, each at a 5 mM final concentration. Although in control assays carried out with yeast that do not express the ABC transporter, we detected a background amount of *t*-R retained in microsomes which was approximately half than that retained in the microsomes of yeast that expressed VvABCB15 (Figure 5B). This transport “blank” is likely due to passive or non-specific endogenous transport as a similar phenomenon has also been reported in other studies such as transport of glycosylated anthocyanidins assayed in yeasts microsomes (Francisco et al., 2013). In the case of *S. marianum* heterologous system, our results show an enhancement in the amount of *t*-R transported of ca. 1.9-fold in cells stably expressing VvABCB15 compared to controls (Figure 6C). Previous studies (Hidalgo et al 2017) showed that *S. marianum* cells expressing grapevine VvSTS3 and incubated with MBCD accumulated extracellular *t*-R, which are the exact conditions we have used as control. It suggests that these cells have endogenous non-specific mechanisms to move *t*-R outwards, and thus the observed enhancement in the double transformants (VvSTS3+VvABCB15) can be attributed to the activity of the transporter. Moreover, the slower disappearance in the extracellular medium of *t*-R externally added to a *S. marianum* cell suspension expressing the transporter provides further evidence that the stable expression of VvABCB15 in this heterologous system gives rise to an active outwards *t*-R transport whether the *t*-R is synthesized in the cell or whether it is taken up from the extracellular medium (Figure 5D).

Here, we carried out two experiments using *Vitis vinifera* cv Gamay cell culture as a homologous system for *t*-R transport functional assays. The rationale behind it relies in its constitutive *t*-R synthesis capacity, which is almost totally accumulated inside the cells in its glycosylated form piceid. In addition, the *t*-R synthesis capacity was handled through the expression of VvSTS3 under control of the light-sensitive promoter pHYH (Gangappa and Botto, 2016) which allowed us to regulate the amount of *t*-R available for transport and glycosylation. Thus, the transport of *t*-R outside the cells promoted by the expression of a candidate gene becomes a competing pathway with the glycosylation and storage that can be conveniently monitored by determining the level of extracellular *t*-R. The stable expression of VvGSTU10 in Gamay grapevine cells gave rise to the accumulation of extracellular *t*-R thus demonstrating its involvement in *t*-R transport out of the cell (Martinez-Marquez et al. 2017). Here we have obtained similar results but carrying out transient expression instead (Figures 7A and 7B), thus validating this type of assay, much less time consuming than stable expression. The effect of transient expression of VvABCB15 in the Gamay cell culture is similar or slightly higher to that of VvGSTU10, and much higher than a mock gene such as GFP in all conditions tested, thus providing strong evidence for the involvement of this particular transporter in *t*-R transport out of cells. On the other hand, the co-expression of both VvGSTU10 and VvABCB15 further increased the transport capacity of the grapevine cells as compared to their individual counterparts (Figures 7A and 7B) pointing towards a cooperative action in *t*-R transport out of grapevine cells. Results obtained in cells expressing both VvABCB15 constitutively and VvSTS1 under control of pHYH and in different light conditions are highly consistent with the above providing strong evidence of the role of VvABCB15 as *t*-R transporter. When comparing the extra- and intracellular profile of stilbenes (Figure 8) it can be noticed that the increase of free stilbene *t*-R in the extracellular medium due to expression of the transporter (no matter whether it is light-inducible or constitutive) corresponds to a concomitant decrease inside of both the free and the glycosylated form piceid. This result can be explained by the competition between the transporter and the glycosylating enzymes for the free *t*-R within the cell and strongly supports the plasma membrane localization of VvABCB15 since if it would localize also in tonoplast the internal *t*-R concentration should have increased as well.

The GST enzymes are long known to be involved in vacuolar accumulation of anthocyanins as well as ABCC type transporters (Goodman et al. 2004), but as the formation of anthocyanin-GSH conjugates is not required for anthocyanin/GSH co-transporters such as VvABCC1 in grapevine (Francisco et al. 2013) or AtABCC2 in Arabidopsis (Behrens et al. 2019) an accepted role of GST is acting as carriers or ligandins to deliver these compounds to the transporters (Sun et al. 2012). Although currently there are no data to support an effective cooperation between these proteins one could speculate with different scenarios. On the one hand, these two proteins might act independently and carry out the transport by parallel mechanism producing additive effects. On the other hand, acting as ligandin as it was proposed for VvGSTU10 (Martinez-Marquez et al. 2017), it would facilitate the movement of the poorly water-soluble *t*-R within the cell and bring it closer to the vicinity of the membrane for transport by VvABCB15, thus enhancing the transport rate by increase of the local *t*-R concentration. Obviously, future experiments that determine for example proximity or even interaction between these proteins are needed to cast light on this issue.

From the work presented herein, it can be concluded that VvABCB15 is a plasma membrane transporter of *t*-R involved in the not yet fully characterized machinery for accumulating *t*-R in the extracellular medium as part of a defense response. To our knowledge, this is the first ABC transporter and the second protein, together VvGSTU10 (Martínez-Márquez et al., 2017), described for this function and future studies will help to elucidate whether other membrane transporters and pumps could be involved in the said machinery. Some other candidates have already been recognized in the proteomics experiment present in this study and by Almagro et al., 2014, and will be investigated in future work.

## MATERIAL AND METHODS

### Plant material

*Vitis vinifera* L. cv. Gamay calli were kindly supplied by Drs. J. C. Pech and A. Latché (ENSA, Toulouse, France) in 1989. This cell line was maintained as both solid and liquid cultures in Gamborg B5 medium as described elsewhere (Bru et al., 2006). *Nicotiana benthamiana* plants were obtained from seeds germinated and grown on potting soil in a greenhouse at a temperature of 25°C, with 16h light/8h dark photoperiod, until they were 3–5 weeks old.

VvSTS3-expressing (Ref. Seq. XM_002263686.2, PREDICTED:stilbene synthase 3 [Vitis vinifera]) *Silybum marianum* transgenic cell line (Hidalgo et al., 2017) was used for stable transformation experiments with VvABCB15. This cell line was maintained as both solid and liquid cultures in MS medium as described elsewhere (Sánchez-Sampedro et al., 2005; Hidalgo et al., 2017).

### Preparation of grapevine cells subcellular fractions and protein extracts

Grapevine cells control and elicited with 50 mM MBCD and 0.1 mM MeJA were harvested after 72h incubation from liquid cell cultures by filtration under gentle vaccum. Subcellular fractions were obtained after mechanical lysis in a potter Elvehjem homogenizer of cell suspensions in 50mM HEPES pH 7.5, 0.25M sucrose,1% (w/v) PVPP, 5% (w/v) glycerol, 10mM EDTA, 10mM Na_2_O_5_S_2_, 10mM ascorbic acid, 1mM PMSF and Sigma Protease inhibitor cocktail at a ratio of 2mL per gram of plant material at 4 °C. Cellular debris was removed by centrifugation at 8000xg for 10min at 4°C and the supernatant (crude extract, Ec) ultracentrifuged at 100,000xg for 1:30h at 4°C. The supernatant (soluble fraction, Fs) was kept apart and the pellet was washed twice by resuspension in double volume of 50mM HEPES pH 7.5, 0.25M sucrose, 5% (w/v) glycerol, 10mM EDTA, 10mM Na_2_O_5_S_2_ and recovered by ultracentrifugation as above. The washed pellet was resuspended in 50mM HEPES pH 7.5, 5% glycerol and layered over a discontinuous sucrose gradient (0, 23 and 32% w/v) prepared in polyallomer tubes and separated by ultracentrifugation at 100,000xg for 3h at 4°C in a swing-out rotor.

Aliquots of the crude extract, the soluble fraction, and each different subcellular sucrose gradient fractions were precipitated as described by Granier and Van of Walle (1988), with slight modifications. Sample was brought to a volume of 750 µl by adding distilled water followed by 8.5 µl of 2% (w/v) sodium deoxycholate solution (Bensadoun and Weinstein, 1975); after mixing and 10 minutes of incubation on ice, 250 µl of 24% (w/v) TCA was is added, vortexed and incubated for 30 minutes on ice to quantitatively precipitate proteins. The protein pellet obtained by centrifugation at 14000xg for 10min at 4°C was washed twice with chilled 10% (w/v) TCA in acetone followed by twice in pure chilled acetone. Finally, the clean protein precipitate obtained was let to dry at room temperature, solubilized in 6M urea and quantified by RC DC protein assay (BIO-RAD) (Raghupathi and Diwan, 1994). One hundred micrograms of precipitated protein sample were digested with trypsin and 30 µg of the resulting peptides desalted with PepClean C-18 Spin Columns (Agilent Technologies) according to manufacturer recommendations as previously described (Martinez-Marquez et al. 2013).

### MRM experiments on protein subcellular markers

The MRM acquisition method was based on surrogate tryptic peptides markers for *Arabidopsis* organelles (Parsons and Heazlewood, 2015; Hooper et al. 2017) suitably adapted to *V. vinifera* protein homologs using Skyline v19.1 software (MacLean et al., 2010). Target marker proteins, their surrogate peptides and transitions for monitoring are schematized in Table S2. MRM analysis were conducted in an Agilent 1290 Infinity LC coupled through an Agilent Jet Stream® interface to a 6490 QQQ triple quadrupole mass spectrometer operated in the positive ion mode. Instrument settings include a spray voltage of 3.0 kV, a nebulizer (psi) of 30, an ion source temperature of 150 °C, gas flow 15 L/min and cell acceleration voltage 4 V. For each transition (table S1), the collision energy applied was optimized in order to detect the greatest possible intensity. The peptide separation was achieved on an Agilent advanceBio Peptide Mapping column (2.1x150mm, 2.7 µm) using a 7 min linear gradient of 3−50% acetonitrile containing 0.1% (v/v) formic acid at a constant flow rate of 0.4 mL/min and an injection volume of 5 µL.

### Label-free proteomic analysis

A proteomic experiment was carried out using triplicates of the tonoplast and the plasma membrane enriched fractions from the grapevine cell suspensions control and treated with 50 mM MBCD+0.1 mM MeJA elicitors for 72 hours. Thirty micrograms of the desalted peptide digests were injected directly in Agilent 1290 Infinity UHPLC coupled through an Agilent Jet Stream® interface to an Agilent 6550 iFunnel Q-TOF mass spectrometer (Agilent Technologies) system. Peptides were separated in a reverse phase Agilent AdvanceBio Peptide mapping column (2.1 mm × 250 mm, 2.7 μm particle size, operated at 50 °C) using a 140 min linear gradient of 3-40 % ACN in 0.1 % formic acid at 0.400 mL/min flow rate. The mass spectrometer was operated in high sensitivity mode. Source parameters employed gas temp (250 °C), drying gas (14 L/min), nebulizer (35 psi), sheath gas temp (250°C), sheath gas flow (11 L/min), capillary voltage (3,500 V), fragmentor (360 V). The data were acquired in positive-ion mode with Agilent MassHunter Workstation Software, LC/MS Data Acquisition B.08.00 (Build 8.00.8058.0) operating in Auto MS/MS mode whereby the 20 most intense ions (charge states, 2 -5) within 300 - 1,700 m/z mass range above a threshold of 1,000 counts were selected for MS/MS analysis. MS/MS spectra (50 - 1,700 m/z) were collected with the quadrupole set to “narrow” resolution and were acquired until 25,000 total counts were collected or for a maximum accumulation time of 333 ms.

Each MS/MS spectra was preprocessed with the extraction tool of Spectrum Mill Proteomics Workbench (Agilent) to obtain a peak list and to improve the spectral quality by merging MS/MS spectra with the same precursor (Δm/z<1.4 Da and chromatographic Δt < 15s). The reduced dataset was searched against the proteome database of PN40024 V2 (CRIBI) genome assembly (Vitulo et al. 2014) and contaminant proteins in the identity mode with the MS/MS search tool of Spectrum Mill Proteomics Workbench and with the following settings: trypsin, up to 2 missed cleavages, carbamidomethylation of Cys as fixed modifications, oxidation of Met, deamidation of Asn and Gln, and pyroGlu as variable modification and mass tolerance of 20ppm for precursor and 50ppm for product ions. Peptide hits were filtered for score ≥ 6 and percent scored peak intensity (%SPI)≥60.

The LC-MS raw files were imported into Progenesis QI for Proteomics (Nonlinear Dynamics) v4.0 label-free analysis software. Quantification was done on the basis of MS1 intensity. The data file that yielded most features (peaks) was used as reference to align the retention time of all other chromatographic runs and to normalize MS feature signal intensity (area under the peak). Correction for experimental variations was done by calculating the robust distribution of all ratios (log(ratio)). The MS features were filtered to include only features with charge state from two to five. After defining the experimental design as “between subjects” mode samples were clustered according to the experimental groups (Control PM, Control T, elicited PM and elicited T) and the average intensity ratios of the matched features across the experimental groups as well as the p-value of one-way ANOVA were automatically calculated. To identify the proteins from which detected features come from, the filtered SpectrumMill peptide hits files were imported into Progenesis QIp, the peptide assignment conflicts resolved (the highest score the winner, or were left unresolved in case of equal score and sequence), and the inferred protein list filtered by score≥15. The abundance of proteins was automatically calculated by the Hi-3 method as described by Silva et al. (2006) implemented in the Progenesis QI for proteomics. Differential protein abundance across experimental group was assessed by using the advanced statistical tools implemented in Progenesis QIp including ANOVA, PCA and Power analysis.

### Functional analysis in yeast microsomes

VvABCB15 transporter (VIT_214s0066g02320; VIT_00032578001; LOC100854950) gene coding sequence codon optimized for *N. benthamiana* was synthesized and supplied inserted into the pESC-URA-cMyc vector (GeneScript; Piscataway NJ, USA). *S. cerevisiae* strain CEN.PK 135 (ura-) was transformed with the pESC-URA-VvABCB15-cMyc vector by the LiAc/SS-DNA/PEG method (Gietz and Woods, 2002).

For in vitro transport studies with yeast microsomes, transformed yeast cells were grown overnight in YNB minimal medium without uracil and spheroplasts were obtained by digestion with lyticase as described (Tommasini et al., 1996). Briefly, yeast cells were cultivated overnight in YNB medium without uracil supplemented with 2% glucose at 30°C. The cells were sedimented at 1200xg for 10 min, washed twice with water, and resuspended in 1.1M sorbitol, 20mM Tris-HCl (pH 7.6), 1 mM DTT containing 57 units of lyticase per ml. After 90 min digestion at 30°C with gentle shaking, the spheroplasts were pelleted by centrifugation for 10 min at 1200xg and lysed in a Dounce tissue homogenizer in 1.1M glycerol, 50Mm Tris-ascorbate pH 7.5, 5mM EDTA, 1mM DTT, 1.5% (w/v) PVP, 2mg/ml bovine serum albumin, 1mM PMSF and Sigma Protease inhibitor cocktail. Unbroken cells and cell debris were removed by centrifugation at 4000 xg for 10 min at 4°C. The supernatants were centrifuged at 100000xg for 45 min at 4°C. Membrane fraction was resuspended in 1.1M glycerol, 50Mm Tris-Mes pH 7.5, 1mM EDTA, 1mM DTT, 2mg/ml bovine serum albumin, 1mM PMSF and Sigma Protease inhibitor cocktail. To study the transport of *t*-R into microsomes, uptake experiments were performed by using the rapid filtration technique (Tommasini et al., 1996.) with nitrocellulose filters (0,45µm pore size, Millipore) as described by Francisco et al. (2013). Briefly, 0.2 ml of yeast microsome suspension (200-350 µg total protein) were mixed with ice-cold transport solution (0.4M glycerol, 100mM KCl, 20mM Tris-HCl (pH 7,4)) and freshly added 1 mM DTT, 5 mM MgSO4, 100µg/mL creatine kinase, and 10 mM creatine –phosphate. A *t*-R and MBCD stock solution in transport buffer was added to give a final concentration of 0.15mM *t*-R and 2.5 mM MBCD, and transport was assayed in the presence of 5 mM Mg ATP and 5 mM GSH in a total reaction volume of 0.8 mL. The transport mixture was incubated at room temperature for 30 min, and then. 0.4 mL of the mixture was immediately loaded on a prewetted filter and rapidly washed with 3 x 5 mL of ice-cold transport buffer supplemented with 20mM MBCD. The filter-bound stilbenes were solubilized by incubating the filter in absolute methanol and extracting for 30 min with shaking. The eluted stilbenes were quantified by MRM as explained below, as described by Hurtado-Gaitan et al. (2017).

### Western blotting

The microsomal pellet was solubilised in 1x SDS-PAGE sample buffer and denatured at 90°C for 5 min. The protein concentration was determined by an RC DC protein assay (BIORAD) (Raghupathi and Diwan, 1994). Proteins (30 µg/lane) were resolved by SDS-PAGE and electro-transferred to the Hybond-P PVDF membranes (GE Healthcare). Membranes were probed at 4◦C overnight with mouse monoclonal anti-Myc-Tag antibodies (Sigma) at the 1:1000 dilution, and were incubated at room temperature for 1 h with horseradish peroxidase-conjugated goat anti-mouse IgG at the 1:10000 dilution. Detection was performed by ECL using the Prime Western Blotting Detection Reagent SuperSignal West Dura system (GE Healthcare, Amersham).

### Cloning of VvABCB15 transporter and VvSTS3 genes

Constructions for constitutive expression of eGFP, VvGSTU10 and VvSTS3 (Figures 6A, 6B and 6C, respectively) are described elsewhere (Martínez-Márquez et al. 2015, 2017; Hidalgo et al. 2017). New DNA constructs for gene expression in plants were designed and assembled using the application GoldenBraid 3.0 standard for modular cloning (https://gbcloning.upv.es/) and cloning parts of the GoldenBraid 2.0 kit (https://www.addgene.org/Diego_Orzaez) (Sarrion-Perdigones et al., 2013). For domestication in the pUPD2 vector, the sequences of interest were PCR amplified flanked by its part category suffix and prefix, and BsmBI recognition sites, as well as removal of internal BsmBI and BsaI recognition sites (See supporting information, Table S2).

Transcriptional units (TU) consisting of a promoter, a coding sequence and a terminator were assembled into pDBG destination plasmids. VvABCB15 TU (pDGB-VvABCB15) consisted of the cauliflower mosaic virus (CaMV) 35S promoter (P35S, GB0030), the VvABCB15 synthetic sequence without the Myc tag and the nopaline synthase terminator (TNos, GB0037) (Figure 4D). In addition, to afford subcellular localization a VvABCB15 C-term fused to YFP TU (pDGB-VvABCB15-YFP) (Fig. 6F) under control of P35S and TNos was assembled. VvSTS3 TU (pDGB-VvSTS3) was built under control of the light responding HY5 HOMOLOG promoter pHYH (Gangappa and Botto, 2016) from grapevine (kindly provided by Tomás Matus) and TNos terminator. A TU for aminoglycoside antibiotic selectable marker was built containing the neomycin phosphotransferase (NPTII, GB0226) coding sequence under control of Pnos and Tnos (Fig. 6D,E).

The GB2.0 α level-assembled transcriptional units (TUs) were further combined within Ω level destination vector backbones, yielding appropriate multigenic constructs as shown in Figure 6. The binary vectors transformed chemically competent *Agrobacterium tumefaciens* strain EHA105 (Hood et al., 1993) by standard techniques (Sambrook et al., 1989).

### Subcellular localization of VvABCB15 in *Nicotiana benthamiana*

*A. tumefaciens* strain carrying the binary vector pDGB-VvABCB15-YFP (Figure 6F) was used for transient expression by agroinfiltration (Schöb et al., 1997) co-infiltrated with a strain bearing cyan fluorescent protein (CFP) as fusion with plasma membrane aquaporin PIP2A (Nelson et al., 2007) for plasma membrane targeting and a strain containing the HC-Pro silencing suppressor (Goytia et al., 2006) in a 1:1:2 ratio in *N. benthamiana* leaves. Three to four days after agroinfiltration the infiltrated leaves were harvested and analysed by confocal microscopy (Leica TCS SP2; Leica Microsystems, Wetzlar, Germany). Leaf segments were cut and abaxial leaf sides were scanned. An argon laser at 514 nm was used to excite the YFP. For visualization, the emission windows were set at 500–545 nm. Serial optical sections were obtained at 1µm intervals, and projections of optical sections were accomplished with the Leica confocal software. Brightness and contrast were adjusted by Adobe Photoshop 7.0. ImageJ with plugin JACoP was used to determine Pearson correlation coefficient.

### Stable transformation of STS-expressing *Silybum marianum* transgenic cells

The *A. tumefaciens* containing pDGB-VvABCB15 (Fig. 6D) was used to stably transform the VvSTS3-expressing *S. marianum* transgenic cell line following the protocol described by Hidalgo et al. (2017). The transformed callus lines were established from individual calli growing in selection medium with 100mg/L paromomycin. Within 2-3 months of the initial transformation, sufficient callus material was obtained to check for plant genome T-DNA integration of the VvABCB15 gene by PCR amplification. The specific primers: Fw 5′-ATGAACTCTTCCTTACAGGTGCC-3′ and Rev 5′-TGAGGTGCCTTGATGATTGC-3′ were used for amplifying a 1097 bp fragment of the VvABCB15 coding region. The amplification reactions are as follows: 1 cycle at 95 °C for 5 min and 30 cycles at 94 °C for 20 s, 54 °C for 30s, 72 °C for 4 min, followed by an extension cycle of 10 min at 72 °C. One transgenic callus was randomly selected and, as shown in Figure 2-suppl the VvABCB15 gene was present in the transgenic line, but not in the parental VvSTS3-expressing *S. marianum* line. One selected transgenic callus material was used to establish rapidly growing cell suspensions.

### Transient transformation of grapevine cells

*A. tumefaciens* harboring constructs (Fig. 6) were used to transiently transform *Vitis* cell suspensions following the protocol described by Martinez-Marquez et al. (2017), but with a 3- or 6-days co-culture and no selection steps. The strains harboring the binary plant vectors pJCV52-VvSTS3 (Hidalgo et al., 2017) (Fig. 6C), pJCV52-VvGSTU10 (Martínez-Márquez et al., 2017) (Fig. 6B) and pDGB-VvSTS3 (Fig. 6E) were used alone or mixed with a strain containing pDGB-VvABCB15 (Fig. 6D) in a 1:1 ratio in *Vitis* cell cultures. From 3 or 6 days after Agrobacterium-infected, stilbenes content was analyzed.

### Elicitor or adsorbent compounds treatments

In Vitis: Treatments were carried out in triplicate as previously described (Bru et al., 2006; Martinez-Esteso et al., 2011a; Martinez-Marquez et al. 2017). Briefly, a weighted amount of filtered and washed cells was transferred into shaking flaks and suspended in fresh growth medium (4mL/g of cell FW) supplemented with either elicitors (5mM MBCD or 50 mM MBCD+0.1 mM MeJA), or adsorbent compounds (1,5 g/L PVP or β-cyclodextrin (βCD)). The cell suspension was incubated with continuous rotary shaking (100 rpm) at 25 °C and under a 16 h light/8 h dark photoperiod.

In Silybum: Treatments were carried out in triplicate as previously described (Sanchez-Sampedro et al., 2005; Prieto D., and Corchete P, 2014; Hidalgo et al 2017). Briefly, 3 g wet weight 14-day old cells were transferred to 100 mL flasks containing 20mL of growth medium and incubated for three days prior to the addition of MBCD-containing medium to 5 mM final concentration. Cultures were incubated in the dark at 25°C and shaken at 90 rpm.

### Determination of stilbenoids

Samples of extracellular and intracellular stilbenes of Vitis cell culture were prepared as described Martínez-Márquez et al. (2016). Then, targeted quantitative analysis of stilbenoids by MRM were performed as described by Hurtado-Gaitan et al. (2017).

Extracellular *t*-R of Sylibum cell culture was extracted three times with two volumes of ethyl acetate as described by Hidalgo et al (2017). *t*-R analysis was performed by HPLC in a Spherisorb ODS-2 (5 μm) reversed-phase column (4.6×250mm) at 35°C. The mobile phase was a mixture of 34 volumes of methanol and 66 volumes of acetic acid:water (5:55 v/v) at 1 mL/min (Hidalgo et al., 2017)). Chromatograms were adquired at 306 nm. Identification of *t*-R was carried out by comparison with a commercial standard and confirmed by LC MS analysis under the same conditions as reported by Hidalgo et al. (2017). Concentrations of *t*-R were estimated using the standard curve generated by pure compound.

## ACKNOWLEDGEMENTS

This work was supported by grants from the Spanish Ministry of Science and Innovation (BIO2017-82374-R and PID2020-113438RB-I00); Valencian Conselleria d’Innovació, Universitats, Ciencia y Societat Digital grant CIAICO/2021/167; European Funds for Regional Development (FEDER) and postdoctoral grant to AMM from Generalitat Valenciana (APOSTD/2018/A/091).

## AUTHOR CONTRIBUTIONS

Conception and design of the work (AM-M, PC and RB-M); acquisition, analysis, or interpretation of data for the work (AM-M, VM, SS-M, HG, PC, and RB-M); writing of the manuscript draft (AM-M, PC and RB-M); all the authors revised and approved the final version to be published, and agreed to be accountable for all aspects of the work in ensuring that any matter regarding the accuracy or integrity of any part of the work are appropriately investigated and resolved.

## SUPPORTING INFORMATION

Additional supporting information may be found in the online version of this article:

**Figure S1:** Label-free proteomic analysis of control (C#) elicited (E#) plasma membrane (M) and tonoplast (T) fractions.

**Figure S2:** HPLC analysis recorded at 306nm of the extracellular medium of transgenic *Silybum marianum* cell cultures stably expressing.

**Table S1:** SRM transitions.

**Table S2:** Oligonucleotides used for GB2.0 domestication and amplification reactions.

**Table S3**: Total proteins identified in label-free proteomic analysis of control (C#) elicited (E#) plasma membrane (M) and tonoplast (T) fractions.

**Table S4:** List of proteins identified with significant differential abundance under the elicitation conditions (ANOVA p< 0.02 and fold change>4) in label-free proteomic analysis of control (C#) elicited (E#) plasma membrane (M) and tonoplast (T) fractions.

**Table S5**: Selected proteins in Figure 2

